# Whole genome error-corrected sequencing for sensitive circulating tumor DNA cancer monitoring

**DOI:** 10.1101/2022.11.17.516904

**Authors:** Alexandre Pellan Cheng, Adam J. Widman, Anushri Arora, Itai Rusinek, William F. Hooper, Rebecca Murray, Daniel Halmos, Theophile Langanay, Giorgio Inghirami, Soren Germer, Melissa Marton, Dina Manaa, Adrienne Helland, Rob Furatero, Jaime McClintock, Lara Winterkorn, Zoe Steinsnyder, Yohyoh Wang, Srinivas Rajagopalan, Asrar I. Alimohamed, Murtaza S. Malbari, Ashish Saxena, Margaret K. Callahan, Dennie T. Frederick, Lavinia Spain, Ariel Jaimovich, Doron Lipson, Samra Turajlic, Michael C. Zody, Nasser K. Altorki, Jedd D. Wolchok, Michael A. Postow, Nicolas Robine, Genevieve Boland, Dan A. Landau

## Abstract

Circulating cell-free DNA (ccfDNA) sequencing for low-burden cancer monitoring is limited by sparsity of circulating tumor DNA (ctDNA), the abundance of genomic material within a plasma sample, and pre-analytical error rates due to library preparation, and sequencing errors. Sequencing costs have historically favored the development of deep targeted sequencing approaches for overcoming sparsity in ctDNA detection, but these techniques are limited by the abundance of ccfDNA in samples, which imposes a ceiling on the maximal depth of coverage in targeted panels. Whole genome sequencing (WGS) is an orthogonal approach to ctDNA detection that can overcome the low abundance of ccfDNA by supplanting sequencing depth with breadth, integrating signal across the entire tumor mutation landscape. However, the higher cost of WGS limits the practical depth of coverage and hinders broad adoption. Lower sequencing costs may thus allow for enhanced ctDNA cancer monitoring via WGS. We therefore applied emerging lower-cost WGS (Ultima Genomics, 1USD/Gb) to plasma samples at ∼120x coverage. Copy number and single nucleotide variation profiles were comparable between matched Ultima and Illumina datasets, and the deeper WGS coverage enabled ctDNA detection at the parts per million range. We further harnessed these lower sequencing costs to implement duplex error-corrected sequencing at the scale of the entire genome, demonstrating a ∼1,500x decrease in errors in the plasma of patient-derived xenograft mouse models, and error rates of ∼10^−7^ in patient plasma samples. We leveraged this highly de-noised plasma WGS to undertake cancer monitoring in the more challenging context of resectable melanoma without matched tumor sequencing. In this context, duplex-corrected WGS allowed us to harness known mutational signature patterns for disease monitoring without matched tumors, paving the way for de novo cancer monitoring.

## INTRODUCTION

Circulating cell-free DNA (ccfDNA) was shown to be a promising clinical tool for non-invasive cancer detection^1–8^. This can be achieved via analysis of cancer-specific epigenetic markers, such as DNA methylation and histone modifications^9–14^. Alternatively, mutation-based approaches using direct genomic sequencing of somatic variants found in circulating tumor DNA (ctDNA)^15–18^ can afford specificity and clinically-actionable information. ctDNA genome sequencing can also aid in detection of low burden of disease, such as cancer screening, detection of minimal residual disease (MRD) after treatment or surgery^17,19–22^, and relapse monitoring^23,24^. In these scenarios, when disease burden is low, the fraction of tumor-derived ccfDNA in the plasma is also low, such that robust detection requires methods with exquisite sensitivity to detect ctDNA signal over the background rate of sequencing error. The technical challenge imparted by the sparsity of ctDNA in low-burden disease settings is typically overcome by increasing sequencing depth at select genomic locations, accompanied by techniques that decrease sequencing error rate. Thus, approaches for reducing sequencing error rates, such as unique molecular identifier (UMI) error suppression techniques^16,25^ or duplex sequencing^18,26–28^, which enable increased accuracy in differentiating true somatic variants from errors introduced by sequencing, can be combined with deep sequencing to optimize successful detection of low-burden disease.

Prevailing methods of ctDNA detection use targeted sequencing protocols, which increase the number of genomes sequenced at a targeted location. However, high throughput targeted sequencing rapidly exhausts available genomes for sequencing (1,000-10,000 genome equivalents (GEs) per mL of plasma^29^), which sets a design-based ceiling on ctDNA detection, where further increases in sequencing depth at targeted sites afford no advantage after the limited number of GEs has already been sequenced. Alternatively, to overcome this limitation, whole genome sequencing (WGS) approaches exploit breadth of coverage to supplant depth, eliminating the reliance on the detection of few sites to increase ctDNA characterization in low tumor fraction settings. For example, our recent method MRDetect^20^ uses primary tumor mutational profiles to inform genome-wide tumor single nucleotide variant (SNV) detection in ccfDNA, such that the available number of GEs is no longer the limiting factor for successful ctDNA detection.

The detection challenges presented by sparsity are significant, calling for broad, accurate and deep ccfDNA sequencing. Thus, whole-genome, low-error, high-coverage methods are necessary for robust ctDNA analysis. However, the costs associated with these approaches are often prohibitive, particularly for clinical application. Although genome sequencing costs have rapidly dropped since the introduction of high-throughput next generation sequencing, more recently this decrease has stagnated^30^. As such, sequencing cost is still a significant barrier for the implementation of high-depth WGS for liquid biopsies, where in clinically important applications, tumor fractions are low (∼10^−5^) and shallow WGS is insufficient for ctDNA detection.

Recently, a new low-cost, high-throughput sequencing method utilizing mostly natural sequencing-by-synthesis (mnSBS)^31^ has been developed by Ultima Genomics. The Ultima platform produces single-end reads at ∼10 billion reads per run for 1USD/Gb, thus substantially lowering sequencing costs compared with current platforms. This cost efficiency holds great promise for many genomics applications, and this approach has now been applied to Genome-In-A-Bottle and 1000 Genomes reference samples^31^ and adapted for single-cell RNA-seq studies^32^. However, to date mnSBS/Ultima sequencing has not been harnessed for application to clinical ccfDNA samples for ctDNA sequencing. In addition, the error rate profiles of this new sequencing method have not been fully characterized, nor have they been rigorously compared with competing technologies. Importantly, for potential application to clinical disease monitoring of ctDNA, it is crucial to have accurate error rate estimates due to the required high sensitivity of ctDNA detection^33^.

Here, to investigate the utility of deep WGS for ctDNA detection, we used the Ultima platform to sequence ccfDNA from plasma samples from healthy controls, cancer patients and patient-derived xenograft mouse models. In a first proof-of-principle study, we show that deep WGS (∼120x) with analytic error correction allows tumor-informed ctDNA detection within the part per million range. We further leveraged the cost-effective and high-throughput nature of mnSBS to develop high coverage WGS duplex error corrected libraries of ccfDNA, achieving error rates as low as 9.2×10^−8^. This allowed us to combine the advantages of genome-wide mutational integration on the one hand, and molecular error correction on the other, to accurately assess disease burden in melanoma patients without the reliance on matched tumor sequencing. Together, our results demonstrate the feasibility and utility of deep WGS for ctDNA detection.

## RESULTS

### Deep sequencing and accurate mutation detection empower low tumor burden detection with plasma whole genome sequencing

Lower limits of ctDNA mutation detection by plasma WGS are dictated by the mutational burden of the tumor, the depth of sequencing, and the rates of error resulting from library preparation and sequencing. To explore these dependencies, we modeled variable tumor fractions (TF), depths of coverages and error rates for a cancer with 10,000 single nucleotide variants (SNVs, approximately 3.7 mutations/Mb) (**Methods**). Tumors with more than 10,000 SNVs are seen across ∼30% of cancers analyzed in the Pan Cancer Analysis of Whole Genomes consortium (PCAWG)^34^ and are enriched in lung (85%), skin (79%), bladder (83%), and other cancers. This analysis suggests that detection of TFs below 10^−5^ requires a depth of sequencing of ∼100x with error rates below 10^−4^ (**Figure 1A**).

**Figure 1.**
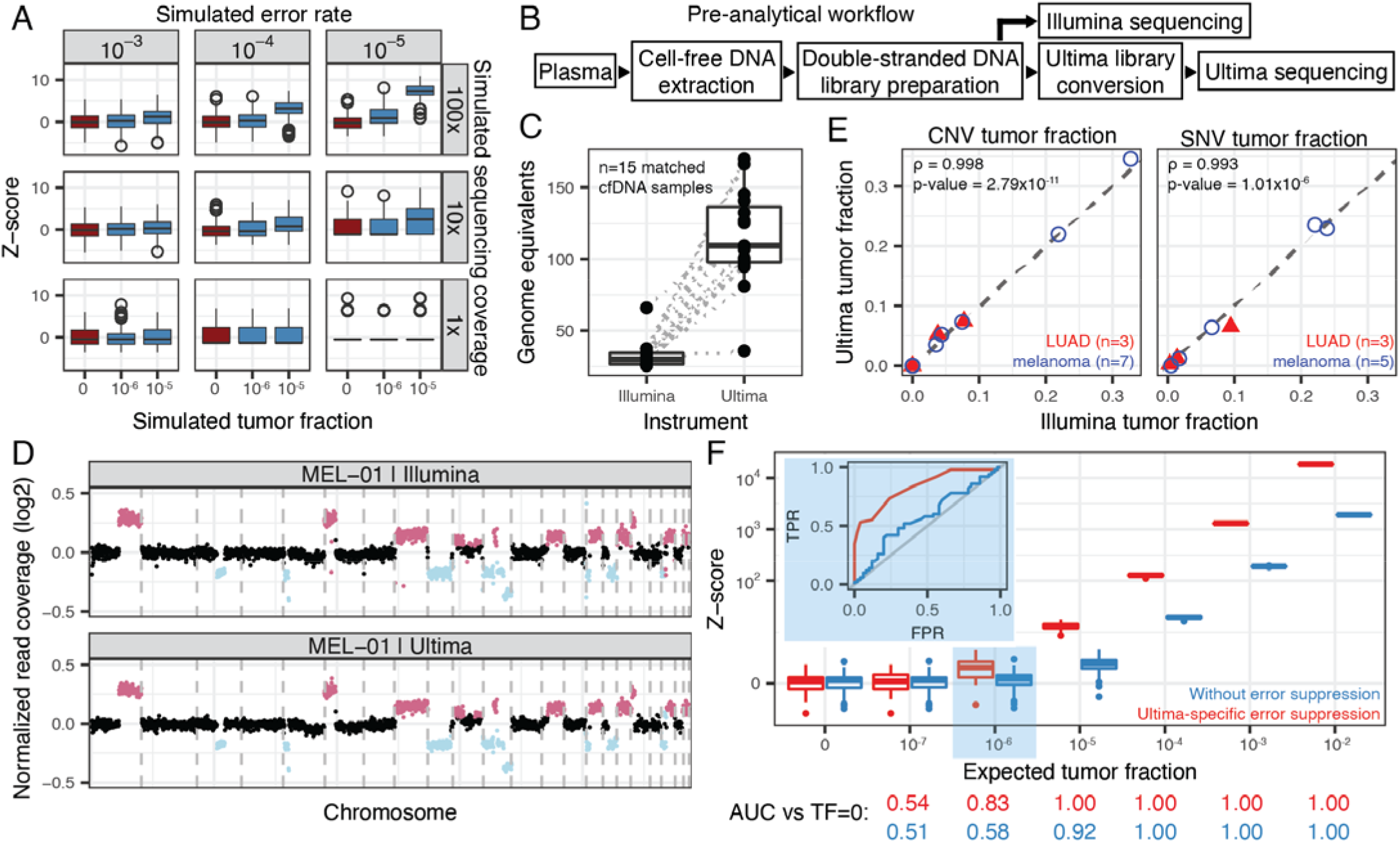
Ultralow ctDNA detection requires deep sequencing coverage and low error rates. A Simulation of ctDNA detection given different error rates (columns), whole genome coverages (rows) and tumor fractions (x-axis). Simulation analysis shows that low error rates and high sequencing coverage are required for accurate ctDNA detection when tumor fractions are at or below 10^−5^ (top right box). B Cell-free DNA library preparation pre-analytical workflow. C Sequencing depth of matched Illumina and Ultima datasets. D Normalized read coverage for Illumina (top) and Ultima sequenced (bottom) matched cell-free DNA sample. E Left: Copy number-based tumor fraction estimation measured with Illumina or Ultima sequencing in matched samples using ichorCNA. Matched cancer-free controls were used to create a panel of normals prior to tumor fraction estimation. Right: Single-nucleotide variant-based tumor fraction estimation measured with Illumina or Ultima sequencing. Somatic SNVs were identified through matched tumor-normal sequencing. Two samples without tumor sequencing and with low ctDNA fraction (<5% measured through CNV analysis) were omitted from this analysis. F In silico mixing study of metastatic melanoma sample MEL-01 with cancer-free control CTRL-05 (50 replicates per tumor fraction, 80x coverage per replicate) with (red) and without (blue) tumor-informed analytic denoising applied using Ultima-specific quality filtering.

Given the need for deeper plasma WGS, sequencing costs impose a significant barrier for broad application. We therefore hypothesized that the recently developed lower cost Ultima sequencing protocol can help overcome this barrier, provided that sequencing error profiles are sufficiently low. To examine this potential application, we performed Ultima WGS on 15 ccfDNA libraries (n = 10 samples from cancer patients [ages 48-86; n = 7 stage IV melanoma and n = 3 III-IV lung adenocarcinoma]; n = 5 controls [ages 58-86, 2 former smokers, 2 current smokers, 1 never smoker]; depth of sequencing of 115x ± 34x [mean ± standard deviation]) with matching Illumina sequencing of the same libraries (33x ± 10x) (**Figure 1B,C, Supplementary Tables 1 and 2**). We first measured circulating tumor DNA burden by estimating the relative abundance of large copy number alterations using ichorCNA^35^ (**Methods**). Tumor fractions had a wide range (<3%, n = 3; 3-10%, n = 5; >10%, n = 2) and measurements were strongly correlated between matched Illumina and Ultima datasets (spearman’s ρ=0.998, p-value = 2.79×10^−11^, Figure 1D,E). Next, we sought to estimate ctDNA burden using a tumor-informed SNV approach. We performed WGS of tumor-derived DNA (or used standard mutation calling on plasma DNA if tumor DNA was unavailable and ctDNA burden was above 5% according to ichorCNA^35^) and matched normal DNA from peripheral blood mononuclear cells (PBMCs) to identify tumor-specific mutations. To remove sequencing errors, we developed a quality-filtering pipeline informed by Ultima-specific feature cutoffs and blacklisted regions (**Supplementary Figure 1, Methods**). Traditional sequencing by synthesis technologies add all 4 bases (A,T,C,G) per sequencing cycle and determine the template nucleotide based on the fluorescence of the complemented base (one base pair is measured per sequencing cycle). However, Ultima’s flow-based technology uses a mix of fluorescently-labelled and unlabeled nucleotides (all nucleotides added in one flow are either A, T, C or G) and relies on the intensity of fluorescence to assign the number of consecutive identical bases that complement the template^32^. Therefore, Ultima technologies may be less susceptible to single-nucleotide substitution errors, given that only a single base type is added per cycle, at the cost of increased difficulty in gauging the size of homopolymers (typically, homopolymers greater than 11 base pairs^31^). Thus, our denoising methods and blacklisted regions were designed to minimize the effect of homopolymer errors instead of single nucleotide substitutions (for example, by ensuring that the 5 flanking bases of a putative variant match the reference, see **Methods)**. We then mined the denoised cell-free DNA reads for somatic variants to estimate ctDNA fractions (see **Methods**) and found strong agreement between Illumina and Ultima sequencing sets (spearman’s ρ=0.993, p-value = 2.01×10^−6^, **Figure 1E**). Next, we intersected ccfDNA reads from cancer-free controls (n = 5) with the SNVs found in patient tumors (n = 6) and measured the frequency of mutations from tumors that were observed in cancer-free control ccfDNA. We found that these error rates were approximately 8.68 × 10^−5^ ± 4.05 × 10^−5^ (mean ± standard deviation, **Supplementary Figure 2**) supporting the utility of Ultima sequencing for detection of low-burden ctDNA.

ccfDNA has been characterized as having a modal length of 167bp with smaller peaks every 10bp, representing the length of DNA wrapped around a single nucleosome and its subsequent degradation^36,37^. Shorter fragment lengths have been associated with ctDNA^38,39^. We therefore compared fragment lengths on matched Ultima and Illumina datasets and found similar distributions below 180 base pairs, underscoring the comparability of the two platforms, with an expected drop beyond 180bp in Ultima datasets (Methods, Supplementary Figure 3).

Finally, to evaluate the impact on the lower limit of detection (LLOD) of tumor-informed, deeply sequenced WGS, we performed an in silico mixing study, as previously shown^20^. We admixed reads from MEL-01 (stage IV melanoma patient, TF = 22%) and CTRL-05 (no known cancer) at different ratios to create admixtures of tumor fractions ranging from 10^−7^-10^−2^ (n = 50 technical replicates per admixture) at 80x sequencing depth. Our group previously developed MRD-EDGE SNV^40^, a cancer-specific deep learning classifier that uses mutation sequence context and other features to analytically distinguish ctDNA from sequencing error at low tumor fractions. Our Ultima-specific denoising framework, in combination with the MRD-EDGE SNV deep-learning architecture, allowed ctDNA detection in the part per million range (Figure 1F), demonstrating the power of deeper WGS to increase ctDNA detection sensitivity. Specifically, performance was evaluated at different simulated tumor fractions using a receiver operating curve (ROC) analysis, showing an area under the curve (AUC) of 0.83 at tumor fractions of 1:10^6^ with analytic error correction, compared to 0.58 in the absence of error correction. These analyses indicate that our framework for ctDNA detection is sensitive enough at low tumor fractions for use in challenging clinical applications such as MRD monitoring.

### Duplex sequencing of ccfDNA applied to the level of the entire genome delivers ultra-low error rates

Recent advances in molecular error correction were shown to radically enhance deep-targeted sequencing approaches, for example through the application of unique molecular identifiers (UMIs) that are incorporated during library preparation for sequencing error correction^16,41^. While strand-agnostic UMIs can help correct for sequencing and PCR errors, UMIs that link forward and reverse DNA strands (i.e. duplex sequencing) can be used to correct for errors that arise on only one strand (such as G>T transversions due to oxidative DNA damage^42^) during library preparation^28^. At the whole genome scale, duplex sequencing was to date cost prohibitive due to the need for high rate of duplicate reads. Nonetheless, studies applying duplex sequencing at the genome scale have shown promise for genome-wide rare variant identification^26,27^. These applications have been limited to ultra-low pass (0.005 GEs/sample^26^) or high input DNA (>50ng^27^) applications, and have not been previously leveraged for liquid biopsies.

We reasoned that the lower sequencing cost afforded by mnSBS can thus be transformative, as it opens the way for affordable genome-scale duplex sequencing in clinical settings. The attendant decrease in sequencing and library preparation errors could be of particular importance for the challenging context of tumor-agnostic (de novo) ctDNA detection, where matched tumor tissue cannot be used to reduce background noise. This advance is of significant clinical importance as the necessity of tumor tissue is an exclusion criterion for many patients^43,44^.

For this important clinical context, we developed duplex WGS for single-end Ultima reads. Here, we created duplex libraries by replacing standard sequencing adapters with adapters containing 3 random nucleotides, thus creating a 6bp duplex-UMI (using the random nucleotides from both ends of the DNA). Libraries were then sequenced using the Ultima sequencer. While duplex sequencing was developed for paired-end sequencing technologies, we were able to recover the ends of most cell-free DNA molecules (80% of all sequenced molecules) due to their modal length of ∼170bp (**Supplementary Figure 3**) to create 6bp UMIs. Then, to ensure compatibility with duplex-processing software that require paired-end reads (here, using the fgbio^45^ suite of tools, **Methods**), we created an in silico paired-end read containing the same mapping and quality information as the original single-end read before processing the alignment files. While duplex error correction offers theoretical sequencing error rates below 10^−9^, true error rates are higher^26,28^ due to mutations mainly originating from library preparation^27^. We note that recent modifications to duplex sequencing offer lower error rates by removing errors resulting from end repair^27^. However, these modifications require either high DNA input for exonuclease use (>50ng) or the use of restriction enzymes, which limit genome coverage. As these requirements are incompatible with ccfDNA WGS, we applied alternative methodology to lower error rates, such as read end trimming (**Methods**).

To first test the accuracy of duplex error correction, we prepared duplex libraries using ccfDNA obtained from the plasma of mice with patient-derived xenografts (n=4, NOD/ShiLtJ species; n = 1 lung cancer; n = 3 diffuse large B cell lymphoma). Tumor fractions, defined as the fraction of reads uniquely mapping to the human genome, were 0.4%, 40%, 73% and 96% (**Supplementary Table 2**). For each sample, we denoised our sequencing reads in three different ways: i) in a UMI-agnostic manner, where PCR duplicates (identified by their mapping positions) are removed from analysis and reads are denoised based on their sequencing and mapping qualities (**Methods**); ii) using UMIs to identify PCR duplicates for single-strand error correction (reads used to create a consensus are profiled base-by-base and a consensus base pair is determined by computing the likelihood of that base being either an A,T,C or G, using the sequenced nucleotides and their qualities as priors); and iii) using PCR duplicates and forward and reverse strands of a same double-stranded DNA template for error correction (duplex error correction). We defined error rates as the number of base pairs in denoised reads that were discordant with the reference genome and occurred two times or less in each position in uncorrected reads (thus removing potential germline mutations), divided by the total number of mapped base pairs in a sample (**Methods)**. Overall, we obtained error rates of 6.8×10^−5^±2.0×10^−5^, 2.2×10^−5^±1.18×10^−5^ and 1.9×10^−7^±1.6×10^−7^ for UMI-agnostic, single-strand corrected and duplex corrected reads, respectively (**Figure 2A**, left), thus achieving a two orders-of-magnitude reduction in error rate with duplex sequencing. When compared to uncorrected reads (without any quality filtering), we observe a ∼1,500x improvement in error rates using duplex WGS (2.0×10^−4^±4.4×10^−5^ error rates in uncorrected reads). Our duplex error rate results are consistent with a previous report employing whole genome duplex sequencing (Abascal et al. reports error rates of 2×10^−7^ using similar protocols^27^), and further suggest that ccfDNA errors are driven more by library preparation and DNA degradation than by sequencing errors.

**Figure 2.**
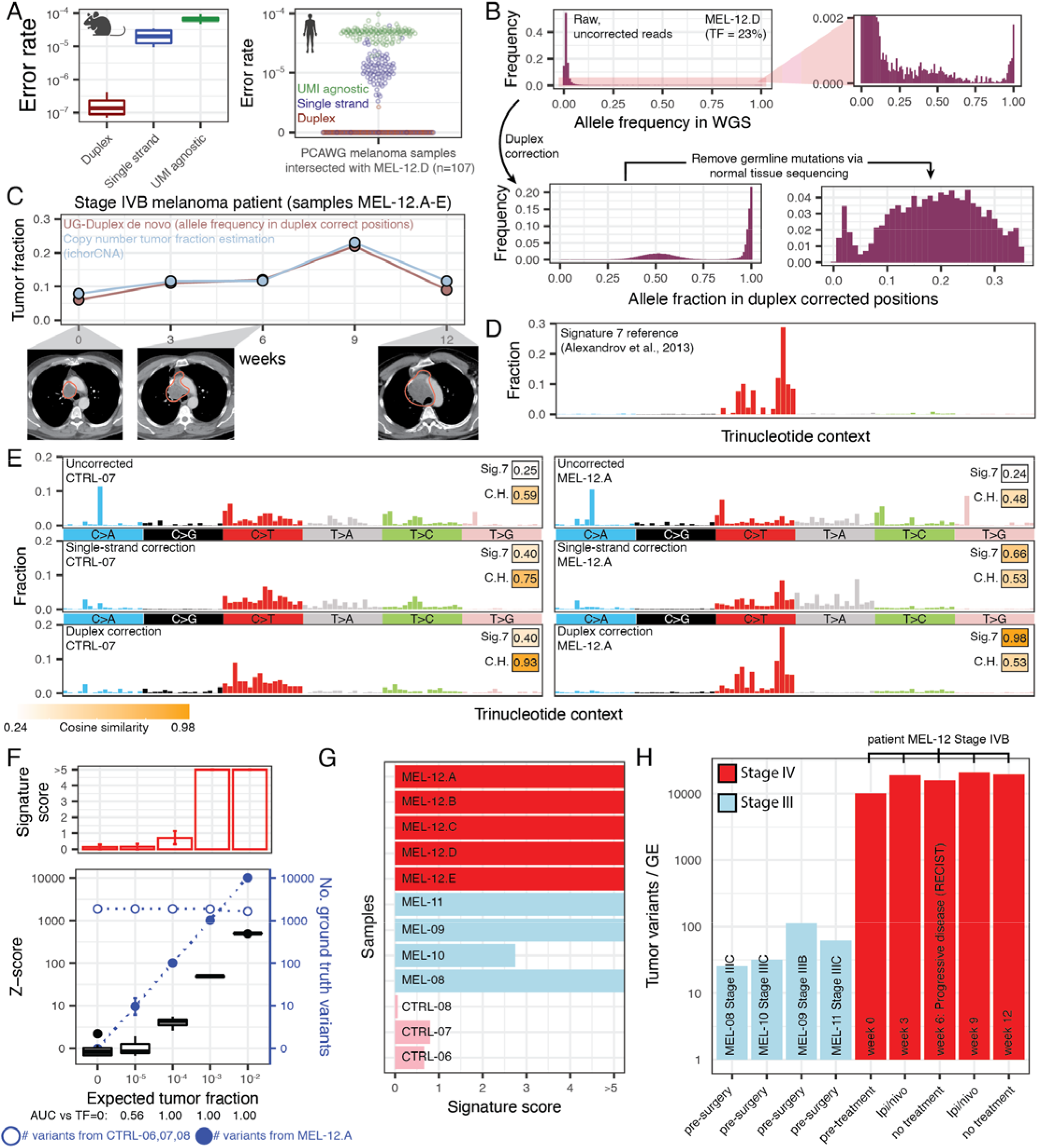
Duplex correction allows ctDNA identification without tumor sequencing. **A** Left: Error rates in WGS sequencing on mouse PDX samples (n = 4). Right: WGS in patient sample MEL-12.D intersected with tumor mutational profiles of 107 melanoma patients retrieved from the Pan Cancer Analysis of Whole Genomes Consortium. Base changes matching the somatic mutation of the tumor were considered errors (after removing germline and somatic mutations from the matched patient data, see Methods). B Variant allele frequencies (calculated using unfiltered sequencing reads) in positions where a variant was found using uncorrected reads (top row) and in duplex corrected reads (bottom row). C Comparison between the modal allele fraction of a patient with progressive disease (samples MEL-12.A-E) in duplex corrected positions (allele fractions above 5% and below 30% only) and copy-number based tumor fraction estimations. D Trinucleotide frequencies from the melanoma-associated UV signature SBS7 reported in Alexandrov et al^47^. E Trinucleotide frequencies in two samples (CTRL-06, left column) and MEL-12.D (right column) in UMI-agnostic corrected WGS, single-stranded correction and duplex correction. Cosine similarity with either SBS7 (UV damage) and clonal hematopoiesis (CH) is compared across conditions. F In silico mixing study of metastatic melanoma samples MEL-12.A with cancer-free control CTRL-06, CTRL-07 and CTRL-08 (10 replicates per tumor fraction, 5x coverage per replicate). Tumor fractions were estimated by fitting the sample’s tri-nucleotide frequencies to that of signatures SBS7 and CH. Top row: signature score to estimate the contribution of signature SBS7 (melanoma UV associated) in the decomposition of a sample’s trinucleotide frequencies into reference signatures. Bottom row: ctDNA detection by expected tumor fraction. Z-scores estimation was used to calculate mutation signature SBS7 detection in comparison to detection in TF=0 replicates. Ground truth variants originating from either the high-burden sample MEL-12.A. or the cancer-free control samples are shown in blue (full circle: MEL-12.A; open circle: cancer-free controls). Error bars represent the standard deviation in the number of variants per replicate at a given expected tumor fraction. G Signature scores of melanoma signature 7 in plasma cell-free DNA samples using duplex WGS (n = 9 melanoma samples; n = 3 controls). Samples in red are from patient MEL-12 with stage IV melanoma at different time points in their clinical course. Samples in blue each represent a separate patient (MEL-08 to MEL-11). Samples in pink represent control samples. H Estimated tumor fraction of samples with elevated signature scores. X-axis depicts the clinical timepoint for each patient sample. Tumor fractions were estimated by multiplying the number of single nucleotide variants found in duplex corrected reads by the weight of signature 7 after reference signature decomposition and normalization by depth of coverage.

To further characterize the accuracy of our duplex sequencing approach, we sought to estimate effective background error rates, that is, the frequency of errors that occur in sites identified in tumors through standard mutation-calling workflows. For this analysis, we performed duplex-corrected plasma WGS on five serial samples obtained from a patient with metastatic melanoma throughout treatment with immunotherapy and obtained duplex coverages of 6.1±3.3 GEs (samples MEL-12A,B,C,D,E). We then intersected somatic variants identified in 107 melanoma tumor mutational profiles from the Pan-Cancer Analysis of Whole Genomes (PCAWG) consortium. As these profiles are unmatched, overlapping mutations may be considered errors. After removing regions with known sequencing artifacts (**Methods**) and filtering for patient specific germline (through matched PBMC sequencing) and somatic mutations (**Methods**), we estimated error rates of 4.7×10^−5^±6.4×10^−6^ (mean ± standard deviation) in UMI-agnostic denoised reads, 9.9×10^−6^±1.8×10^−6^ in UMI single-stranded corrected reads, and 1.4×10^−7^±4.0×10^−8^ in duplex corrected reads (**Figure 2A**, right). We note that these error rates likely represent the upper bound of the true error rate, as rare variants from diverse origins such as clonal hematopoiesis^46^, sample contamination^27^ or somatic mosaicism^27^ would be classified as errors here. Collectively, these analyses demonstrate that duplex error correction applied to the scale of a whole genome results in ultra-low error rates. To further illustrate the potential of duplex error correction for variant calling in ccfDNA, we first assessed the variant allele frequency (VAF) distribution in non-cancer control ccfDNA. While UMI-agnostic WGS showed only 75%±4.8% of base substitutions identified at the expected VAF range for germline events (see Methods and Supplemental Figure 4), this increased to 99%±0.0005% after duplex error correction. In high burden metastatic melanoma ccfDNA samples, the application of duplex error correction and removal of germline mutations allowed the emergence of somatic mutation VAF modes consistent with tumor fraction estimates using ichorCNA (**Figure 2B,C**), rendering this approach feasible for ctDNA mutation detection in clinical samples without a matched tumor (non tumor-informed).

### De novo (non tumor-informed) monitoring of low burden cancer through mutational signature patterns in duplex ccfDNA WGS

Current methods for de novo detection of somatic mutations (i.e., non tumor-informed) rely on “off the shelf” sequencing panels that target recurrent mutations of a given cancer-type. These methods are inherently limited by the scarcity of cell-free DNA in plasma (only up to 1,000-10,000 genomic equivalents in a mL of plasma^29^) and require a given patient’s cancer to harbor the targeted mutations. In addition, somatic mutations in ccfDNA can also originate from mutations occurring in blood cells rather than ctDNA of solid tumor origin. For example, previous studies have identified mutations arising from clonal hematopoiesis as significant background, hindering ctDNA detection^48^.

WGS-based non tumor-informed mutation detection requires a different perspective as the detection of individual cancer hotspots in WGS is not sensitive due to relatively lower coverage per genomic position. Thus, SNV-based WGS methods of ctDNA detection must rely on genome-wide mutational integration, where the identification of multiple mutations improves detection power^20^. We hypothesized that signatures of somatic mutation accumulation^46,49,50^ could serve as such a framework for genome-wide mutational integration for non tumor-informed WGS ctDNA detection.

Specifically, we reasoned that genome-wide mutations from the sequencing reads could be integrated and summarized as a weighted sum of single-base substitution (SBS) reference mutational signatures and could originate from the tumor in cancer patients and clonal hematopoiesis (CH)^46^. We explored the trinucleotide contexts of ccfDNA variants through mutation signature analysis at each level of denoising (UMI-agnostic, UMI single-stranded and duplex), to investigate the potential sources conferring alterations in these samples. Cosine similarities (often used^49^ to measure the similarities between two mutational signatures) between the UV-associated SBS7 signature (COSMIC^51^ v2) and high burden samples (MEL-12A-E, stage IVB melanoma) were highest after duplex correction (mean cosine similarities to SBS7 of 0.38±1.58 (range 0.24-0.64), 0.90±0.15 (0.66-0.99), and 0.99±0.006 (0.98-0.99) between UMI-agnostic denoising, single-strand and duplex corrected datasets, respectively, **Figure 2D,E** and **Supplementary Figure 5**). We found similar improvements when measuring cosine similarities between the clonal hematopoiesis signature and cancer-free controls, highlighting the importance of duplex correction for accurate signature analysis, and demonstrating that clonal hematopoiesis is an abundant source of mutations in ccfDNA WGS (**Figure 2E, Supplementary Figure 5**).

Given the ability of de novo mutation identification in error corrected ccfDNA WGS to deliver profiles matching SBS7 and CH signatures for identifying melanoma and age-associated circulating DNA fragments, respectively, we developed a tumor-agnostic approach for ctDNA detection based on mutational patterns. As a first step, we tabulate the tri-nucleotide frequencies of plasma ccfDNA mutations, which are fitted to the reference mutational signatures (here, SBS7 and CH) using a non-negative maximum likelihood model. The relative contributions of the reference mutational signatures are thus obtained, and a tumor score is estimated by taking the weight of the tumor-associated SBS7 signature and multiplying by the number of variants per duplex genomic equivalents sequenced. Then, a signature score is calculated to empirically determine the importance of SBS7 to the estimated tumor score. The signature score is obtained by repeating the fit of the plasma ccfDNA mutations to the reference, but the tri-nucleotide frequencies of SBS7 are randomly permuted N times for each fit. We then count the number of times (P) the weight of SBS7 is increased after permutation and calculate the signature score as -log(P / N).

To analytically validate our approach, we performed an in silico mixing study combining duplex-denoised reads from a high burden ctDNA samples (MEL-12.A, tumor fraction estimate of 7.9% via ichorCNA; 148,819 variants at 9.3x duplex sequencing coverage) and the three cancer-free controls (CTRL-06, CTRL-07, CTL-08; 2,936 variants at 7.77x aggregated duplex sequencing coverage) at 5x duplex sequencing depth, in varying proportions (expected tumor fractions of 0, 10^−5^-10^−2^). We found that signature scores were high (greater than 5) at tumor fractions greater or equal than 10^−3^. Signature scores dropped at tumor fractions at or below 10^−4^, given that the number of mutations originating from ctDNA was significantly below the number of mutations arising from cancer-free ccfDNA (∼100 vs 2,000 variants at 10^−4^, from MEL-12.A vs from cancer-free controls, respectively, **Figure 2F**). However, estimated ctDNA was still readily detectable at tumor fractions of 10^−4^ (receiving operating characteristic area under the curve 1.00 compared to TF=0) (**Figure 2F**). Tumor scores correlated strongly with expected tumor fractions (spearman’s ρ = 0.88, p-value <2.2×10^−16^). Importantly, these results highlight that we can identify ctDNA contributions when the number of tumor-derived variants is below the number of variants originating from background biological processes such as CH (∼100 cancer-derived mutations in ∼2,000 background mutations, at tumor fraction of 10^−4^), which are expected to dominate rare variant signal in early-stage and minimal residual disease contexts.

Next, we applied our signature-based ctDNA detection platform for pre-operative ctDNA detection (i.e. tumor-agnostic ctDNA detection). We sequenced plasma samples from four patients with resectable locoregional stage III melanoma (without tumor or normal DNA available), three cancer-free controls, and one treatment-unresponsive patient with stage IV melanoma (5 separate time points, patient MEL-12; with PBMC-derived normal DNA available). Detection of locoregional melanoma is clinically relevant, given that stage IIIA has a 5 year survival rate of 93%^52^. However, current methods often fail to detect somatic variants in plasma of stage III melanoma patients^52,53^. In our cohort, tumor fractions were readily detectable via ichorCNA for the stage IV melanoma samples (**Figure 2C**) but were undetectable in samples from patients with stage III disease, suggesting tumor fractions below 3%, the limit of detection of ichorCNA^35^ (**Supplementary Figure 6**). Denoised duplex reads from stage III samples were kept if they were the only read carrying a given variant, thereby reducing the probability of germline mutations. We note that this form of germline variant filtering is not expected to result in substantial loss of ctDNA, given that the tumor fractions are below 3%, and thus we could reasonably expect, at most, one supporting ctDNA read at any given tumor-derived mutated locus.

Tumor scores were readily separable between the control, stage III pre-operative melanoma and stage IV melanoma samples (3.66±4.06, 58.03±39.65 and 17,104.10±4,304.55, respectively, Figure 2H). Previous studies have shown that undetectable ctDNA using targeted panels is associated with favorable prognosis in stage III melanomas. However, patients with undetectable ctDNA often still experience disease recurrence, highlighting the need for more sensitive tools for improved stratification. Here, signature and tumor scores for ctDNA detection showed strong separation between cancer-free controls and samples from melanoma patients (**Figure 2G-H**). Our results thus suggest that deep error corrected sequencing can identify ctDNA in a non tumor-informed approach using mutational signatures, even when ctDNA burden in the plasma is low such as in resectable melanoma.

## DISCUSSION

High-throughput short-read sequencing has revolutionized the liquid biopsy field, and sequencing costs have historically decreased at a rate faster than Moore’s Law^30^. However, the decrease in cost has stagnated over the last 5 years. While sequencing costs are less of a barrier for targeted panels given that they profile < 2% of the genome, targeted panel sensitivity is limited by the low abundance of cell-free DNA in the plasma. Conversely, WGS overcomes the ccfDNA abundance barrier by profiling the entirety of the genome, but is limited by costs of sequencing.

Ultima Genomics has recently debuted a mostly natural sequencing-by-synthesis platform that generates low-cost (1USD/Gb) whole genome sequencing^31,32^. To date, Ultima sequencing has been benchmarked in germline variant calling^31^ and single-cell RNA sequencing^32^. While these data are informative for these specific experimental contexts, several important technical performance parameters in the context of deep ccfDNA WGS remained unknown. Specifically, it was unclear if Ultima sequencing could operate at sufficiently low error rates to be used for ctDNA detection, where error rate is critical, or if the technology could be leveraged for deep WGS or duplex sequencing, neither of which had been explored previously. Thus, to assess Ultima’s suitability for low-error, deep WGS applications, here, we performed a comparative analysis of Ultima and Illumina short-read sequencing platforms, generating comprehensive error rate profiles and demonstrating that these two approaches have comparable tumor-informed analysis capabilities for circulating tumor DNA detection (**Figure 1**). Our analyses demonstrated that Ultima sequencing is suitable for deep WGS and provided direct comparison to ccfDNA with Illumina WGS, paving the way for broader application and adoption by the genomics community.

Importantly, we find that higher depth of coverage in WGS tumor-informed approaches allows for ctDNA detection even at the parts per million range, thereby making such analysis suitable for minimal residual disease detection. In such scenarios, where ctDNA frequencies are well below 10^−4^, targeted sequencing is often insufficient for ctDNA detection due to the limited number of genome equivalents, and further increasing sequencing depth affords no advantage. In contrast, our modeling shows that WGS approaches for ctDNA detection benefit from higher sequencing depth, where detection of tumor fractions below 10^−5^ is possible by accurate (error rates below 10^−4^), deep (∼100X) sequencing (Figure 1A). Hence, the cost-efficiency of the Ultima platform can be harnessed for ctDNA detection by high-depth WGS at tumor fractions reflective of minimal residual disease (MRD), where deep panel sequencing has been shown to be inadequate (e.g., in only 20–70% of cases positive for early-stage cancer by imaging had ctDNA detected by deep panel sequencing^15,21,54^).

Given that we can achieve ctDNA detection rates in the parts per million range with high-depth WGS, to further expand and demonstrate the utility of the technology, we addressed the more challenging problem of ctDNA detection in tumor-agnostic (non tumor-informed) settings. We developed single-end whole genome duplex sequencing and computational denoising (**Methods**) to achieve low error rates (∼10^−7^) and deconvolve the cell-free DNA mutational compendium into representative mutational signatures to detect ctDNA without matched tumor sequencing (**Figure 2**). Specifically, we leveraged the fact that somatic single nucleotide substitutions display characteristic enrichment in particular nucleotide contexts, and that these patterns have been defined as mutation signatures of known (or sometimes unknown) sources^46,49,50^, for example UV signature in melanoma. In this way, the genome-wide SNVs from ccfDNA can be collapsed into mutation signatures, compared with the established reference signatures, and assigned as being of cancer origin or not, without the need for a comparator tumor sample with genotyped SNVs. Thus, this tumor-agnostic, mutation signature-informed approach represents a powerful way to untether ctDNA detection from patient-specific sequencing panels.

Compared to commonly-used off-the-shelf panels, whole-genome analysis has the benefit of sequencing breadth, allowing for the detection of tumor-derived mutations that may not be present in targeted panels. In addition, whole-genome analysis enables analytical distinction between ctDNA-derived mutations and those arising during clonal hematopoiesis, which can confound ctDNA detection^48^ (**Fig. 2**). Finally, as highlighted above, the breadth of coverage in whole-genome analysis removes the sequencing depth ceiling imposed by limited number of GEs for sequencing at low tumor fractions. While larger patient cohorts and applications to different cancer types are necessary to validate our cancer-detection platform, the work presented herein is a proof-of-principle study that shows the potential of deep error-corrected WGS for sensitive ctDNA detection. We anticipate that in addition to melanoma, this approach will be applicable to other cancer types where there are a high number of SNVs carrying a distinct mutational signature, such as smoking-associated lung cancer^47,49^. We envision that our methods can thus be harnessed for de novo cancer monitoring in low burden disease scenarios, providing a powerful tool for diagnosing cancer and detecting relapses at the earliest stages.

## CONCLUSION

Radical decreases in sequencing costs open new opportunities in clinical genomics. Here we harnessed low-cost mnSBS to demonstrate that impact of deeper WGS (∼100X) on tumor-informed ctDNA monitoring ctDNA. Moreover, we also developed methodology to integrate duplex sequencing with the single-end mnSBS. This allowed to apply duplex error correction to plasma clinical samples at the level of the entire genome. We leveraged these highly error-corrected data for a non tumor-informed ctDNA detection, based on the similarity of mutational profiles to known cancer mutational signatures. These findings have important clinical implications, as uncoupling ctDNA detection from a tumor-informed mutation profiles radically increases the potential use of ctDNA monitoring in common clinical scenarios where tumor samples cannot be obtained. Excitingly, this demonstration opens up the possibility for the use of WGS in ctDNA cancer screening. In particular, such an approach might be beneficial for specific settings where there is high genetic (e.g. BRCA mutation carriers, Lynch syndrome) or environmental (e.g. tobacco smoke exposure) cancer risk together with distinct mutational signatures.

## ACKNOWLEDGEMENTS

We thank the patients and their families for contributing plasma and tissue for this study. We also thank members of the Landau laboratory, the New York Genome Center computational biology team, especially Minita Shah, and the New York Genome Center research sequencing laboratory for thoughtful discussions throughout this work. We thank Dr. Catherine Potenski for critical evaluation of the manuscript. This work was supported by the Mark foundation Aspire Award, the Burroughs Wellcome Fund Career Award for Medical Scientists, National Institutes of Health Director’s New Innovator Award (DP2-CA239065) and a National Cancer Institute R01 grant (R01-CA266619-01) (DAL). MSKCC investigators are supported by Cancer Center Support Grant P30 CA08748 from the National Institutes of Health/National Cancer Institute. A.J.W. received support from the Conquer Cancer Foundation Young Investigator Award. The opinions, results, and conclusions reported in this paper are those of the authors and are independent from these funding sources.

## CODE AND DATA AVAILABILITY

The raw genomic sequencing data and clinical variables are being deposited to the European Genome-Phenome Archive. Code and custom scripts will be made available as a GitHub repository.

## Methods

### Simulation analysis

Simulations for ctDNA detection scores (Figure 1A) were performed assuming a tumor-mutational compendium of 10,000 SNVs with different error rates (10^−3^, 10^−4^ and 10^−5^), coverages (1, 10 and 100) and tumor fractions (0, 10^−6^, 10^−5^). For each of the 10,000 SNV mutations, coverage was simulated using a Poisson distribution^55^. Each simulated sequenced base pair was classified as either ctDNA or ccfDNA according to the tumor fraction, and errors misclassified as ctDNA were determined according to the error rate. Estimated tumor fractions were calculated by summing the ctDNA molecules and the errors and dividing by the total base pairs simulated. Z scores were calculated as:

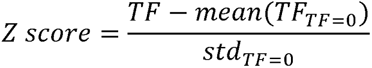

### Tumor and germline DNA extraction, library preparation and sequencing

Genomic DNA was extracted using the QiAamp DNA Mini Kit (Qiagen, cat#56304) and the QIAamp DNA blood Kit (Qiagen, cat#51104) for tissue and blood samples, respectively, and sheared to 450bp (Covaris cat#500569). Sequencing libraries were prepared on 1µg of DNA using the TruSeq DNA PCR-Free Library Preparation Kit (Illumina, cat#20015963), with one additional bead cleanup performed after end-repair and after adapter ligation. DNA was quantified using a Qubit 3.0 fluorometer and length analysis was performed using an Agilent Bioanalyzer or High Sensitivity Fragment Analyzer. 2×150bp paired-end sequencing was performed on either a HiSeq X or NovaSeq v1.0 Illumina machine.

### Pan-Cancer Analysis of Whole Genomes (PCAWG) datasets

We downloaded n = 107 datasets from the PCAWG database (Supplementary Table 3). These datasets correspond to somatic single nucleotide variants from whole genome sequencing of melanoma tumor tissue. Specifically, these datasets originate from the SKCM-US and MELA-AU cohorts, processed through the PCAWG SNV-MVN calling pipeline.

### Human subjects sample processing

Blood samples were obtained from patients after obtaining informed consent and following protocols approved by institutional review boards and in accordance with the Declaration of Helsinki protocol. Samples were obtained from either NewYork-Presbyterian/Weill Cornell Medical Center, Memorial Sloan Kettering Cancer Center, Massachusetts General Hospital, or the Royal Marsden NHS Foundation Trust in the United Kingdom (Supplementary Table 1). Tumor, normal and plasma samples from the Royal Marsden NHS Foundation Trust were obtained under an ethically approved protocol (Melanoma TRACERx, Research Ethics Committee Reference 11/LO/0003). Tumor tissues were collected from resected lung or melanoma specimens. Cancer diagnosis was established according to World Health Organization criteria and confirmed in all cases by an independent pathology review.

### Cell-free DNA extraction

Cell-free DNA was extracted from plasma using the Magbind ccfDNA extraction kit (Omega Biotek cat#M3298). Manufacturer recommendations for extraction were followed, but elution volume was increased to 35 µL, and elution time was increased to 20 minutes on a thermomixer at 1,600 rpm (room temperature). Extracted ccfDNA was quantified using a Qubit 3.0 fluorometer and length analysis was performed using an Agilent Bioanalyzer or High Sensitivity Fragment Analyzer.

### Cell-free DNA library preparation

Whole genome library preparation (without duplex) Next generation sequencing libraries were generated using a double-stranded preparation kit (Kapa Hyper Prep Kit, Roche, cat#KK8502). Full-length adapters (IDT TruSeQ UDI plate, Illumina, cat# 20023784) were used for adapter ligation. Six PCR cycles were carried out when input DNA was above 5ng, and 8 cycles were performed when the input was below 5ng. Libraries were quantified using Qubit 3.0 fluorometer and length analysis was performed using an Agilent Bioanalyzer or High Sensitivity Fragment Analyzer. Illumina sequencing libraries were sequenced on a HiSeq X or NovaSeq1.0 using 2×150 paired-end sequencing.

#### Whole genome library preparation (with duplex)

ccfDNA libraries were generated in a similar fashion as described above, although the full-length adapters were replaced with stubby Y-adapters containing 3 UMI bases (IDT Duplex Seq adapters, cat#1080799) and sample indexing was carried out during PCR amplification. To enhance duplicate recovery in human samples, 4ng of prepared libraries was subjected to 6 additional PCR cycles prior to Ultima library conversion. Mouse PDX samples did not undergo additional PCR cycles prior to Ultima library conversion.

#### Ultima sequencing

Illumina sequencing libraries underwent Ultima library conversion. Briefly, Illumina libraries were converted to Ultima libraries by PCR, using primers matching Illumina read 1 and read 2 sequences and containing Ultima-specific barcodes (R1 conversion adapter: TCC ATC TCA TCC CTG CGT GTC TCC GAC TGC ACA ATG TGT GCT AGA TCT ACA CGA CGC TCT TCC GAT CT, R2 conversion adapter: CTG TGT GCC TTG GCA GTC TCA GCT CAG ACG TGT GCT CTT CCG ATC T). Samples were then pooled and sequenced on an Ultima sequencer prototype.

### Whole genome sequencing (without duplex) adapter trimming and alignment

Illumina fastQ reads were adapter trimmed using skewer^56^ (version 0.2.2). Trimmed reads were then aligned to the human genome (version hg38) using bwa mem. Duplicate reads were marked in a UMI-unaware fashion using novosort^57^. Depth of coverage was estimated using mosdepth^58^ (version 0.2.9). For Ultima reads, adapter trimming was carried out to remove the Illumina conversion adapters. Cutadapt^59^ (version 2.10; cutadapt --mask-adapter -a CTACACGACGCTCTTCCGATCT;max_error_rate=0.15;mi n_overlap=10;required… AGATCGGAAGAGCACACGTCTGCTG;max_error_rate=0.2 ;min_overlap=6) was used to mask adapter sequences, and adapter trimming was then carried out using GATK^60^ (private Ultima fork, since merged to the latest 4.3.0.0 GATK release) (ClipReads function). Alignment was performed using bwa mem^61^ (version 0.7.15-r1140).

### Copy-number based tumor fraction estimation

Genome-wide coverage was calculated over a 1Mbp window and normalized for mappability and GC content biases (using hmmcopy^62^ version 0.99). Tumor fractions were estimated using ichorCNA^35^ (version 0.3.2) after correcting for library and sequencing artifacts via a panel of normals (PoN) from cancer-free controls (CTRL-01 to CTRL-05). A separate PoN was created for Illumina and Ultima sequenced samples using libraries sequenced on the respective machines. For plotting purposes (Figure 1D), corrected log2 read counts outputted by ichorCNA were used. Bins marked by ichorCNA as copy gains, amplifications and high-level amplifications were marked and colored as chromosome gains (pink). Bins marked as homozygous deletion states and hemizygous deletions were marked and colored as chromosome losses (blue). Copy neutral regions were marked as neutral (black). Bins with corrected log2 read counts between -0.05 and 0.05 were marked as neutral (black) as well.

### Whole genome sequencing (without duplex) SNV-based tumor fraction estimation

SNV-based tumor fraction estimation was carried out by counting cell-free DNA reads with matching tumor-specific somatic mutations. To limit the effect of problematic regions of the genome, a platform-specific blacklist was built. For Illumina sequencing, regions identified in the ENCODE blacklist^63^, centromeres^64^, simple repeat regions^64^ and positions with high mutation rates (GNOMAD^65^, AF>0.001) were not considered. For Ultima sequencing, Ultima-specific low-confidence regions composed of homopolymers, AT-rich regions, tandem repeats, and regions with poor mappability and high coverage variability were additionally excluded (Supplementary figure 1).

To limit the effect of sequencing errors, custom scripts were used for platform-specific denoising. Illumina alignment files were filtered to contain read pairs overlapping somatic mutation positions. Paired-end reads were filtered for mean base pair quality (greater or equal to 10), base pair quality of the queried position (greater or equal to 25), template length (below 240), and coverage of the position (greater or equal to 25) and were only kept if both R1 and R2 carried the mutation. Tumor fractions were estimated by dividing the number of filtered reads containing the somatic mutation by the total number of filtered reads.

Ultima alignment files were subset to reads overlapping with somatic mutation positions (bedtools^66^, version 2.29). Reads were filtered by mapping quality (mapping qualities below 60 were discarded) and template length (measured as the length of the sequencing read after adapter trimming; reads greater than 200bp were discarded). Variants were identified using the FlowFeatureMapper tool from the UG-specific GATK suite of tools with additional filters. Reads were only considered if the five bases flanking each side of a variant were identical to the reference, and if the flow-based cycle score was greater or equal than 3. Ultima sequencing data is flow-based in nature^31^, so that in each sequencing cycle a single nucleotide base type is incorporated, and the length of the respective homopolymer is measured. Sequencing errors are therefore most commonly homopolymer indels, while substitution error can occur due to changes in one or more flow cycles. In order to assign qualities, or sequencing error likelihoods, to substitutions, Ultima has developed the FlowFeatureMapper tool that was made public as part of the Genome Analysis Toolkit 4.3.0.0 release. Briefly, the tool extracts all the single nucleotide substitution from a CRAM file, filters them and assigns a likelihood score. In this study, reads were filtered for MapQ=60, and each substitution was required to match the reference in the adjacent -+5 base pairs. The former was done in order to avoid alignment artifacts, and the latter to avoid indels being interpreted as substitutions by the aligner. A likelihood score is then calculated by considering two candidate short local haplotypes – one matching the reference genome and one differing in the considered substitution. Using the flow-homopolymer probability matrix decoded from the UG CRAM files, describing the probability for each homopolymer in each flow cycle^31^, the log likelihood of the read to have arisen from each of the two haplotypes is calculated, and their difference is reported as a log likelihood score (X_SCORE). This score can be interpreted as the likelihood of a substitution to be the result of a sequencing error. Substitution were filtered for a minimal quality 3, corresponding to an error likelihood of 0.1%.

Tumor fractions were estimated by dividing the number of filtered reads containing the somatic mutation by the total number of filtered reads. For the lower limit of detection estimation, denoised-reads were processed through MRD-EDGE^40^ as described below.

### SNV model training sets and feature space

Our training sets were obtained from plasma enriched for ctDNA SNV fragments (true label) from specific melanoma tumors and ccfDNA SNV reads from Ultima datasets (false label) from healthy controls without known cancer. Candidate reads were extracted from our custom denoised alignment files. For our true label sets we used patients with high burden metastatic disease and only reads which represented matched tumor variants were retained. We used a custom deep-learning model for signal to noise enhancement^40^ to categorize candidate SNV reads. Briefly, an ensemble model using CNN and MLP classifiers was used.

#### CNN model architecture

A one-hot encoded tensor structure was used for each candidate read containing a single-nucleotide variant. The encoded tensor has and image-like structure with a shape of 12×240. The rows correspond to one hot encoded nucleotides (N, A, C, T, G) corresponding to the reference and the read. The penultimate row dimension is used to mark the position along the read highlighting the SNV of interest. Lastly, a binary version of the variant flow score (1 = flow score of 10; 0 = flow score below 10) is encoded along the last row dimension to add further relevance to tri-nucleotide context of the SNV of interest. The columns correspond to individual nucleotides along the length of the read. While our reads are mostly below 200 base pairs (Supplementary Figure 1), the extra base pairs are padded with the reference genome to add additional contextual information. Estimated error rates were obtained by running the cancer-free controls through the aforementioned pipeline. Each control was intersected with the mutational landscape of each tumor. The rate of denoised reads matching the somatic mutations of a tumor was taken as the error rate.

#### MLP model architecture

A tabular set of feature values is provided as an input to the MLP. Epigenetic features (available at Supplementary Table 4) at potential variant positions as well as read-specific quality metrics were used to distinguish true variants from sequencing noise. Read quality metrics included Levenshtein edit distances of the read upstream and downstream of the variant, forward or reverse mapping strands of the read, the number of SNVs present on a read, as well as the number of SNVs present on a read where the 5 flanking bases match the reference genome.

### SNV model design and training

Our deep-learning model has an ensemble structure and consists of two major components -a regional/read specific multi-layer perceptron (MLP) and a sequence based convolutional neural network (CNN), whose weight matrices are jointly learnt. The MLP which takes a feature matrix as input consists of a linear stack of four dense blocks. We define each block as consisting of a fully connected layer with a ReLU activation. Furthermore, for the purpose of regularization the input to each fully connected layer is batch normalized and the output is passed through a dropout layer. The CNN consists of four one dimensional convolution layers with non-linear ReLU activations, which extract sequential information at different spatial resolutions. Moreover, as in classical deep learning frameworks, each convolution layer (post nonlinear activation) is followed by a max pooling layer. The output is then passed through a stack of 3 dense blocks as defined above. Subsequently, the latent output of both the MLP and CNN is then concatenated and passed through a single dense block. Finally, a probability score between 0 (sequencing noise) and 1 (true somatic mutation) is obtained by using a single sigmoid-activated fully-connected layer. This probability score reflects the model’s estimate on whether a candidate SNV mutation present in the encoded read is likely from signal or noise. Our ensemble model is built in Keras (v.2.3.0) with a Tensorflow base (1.14.0). To train our ensemble model, we minimize the objective function defined as a binary cross entropy loss. We report our performance metrics within balanced sets.

### Analytical lower limit of detection estimation

We created in silico mixes at various tumor fractions by computationally combining aligned reads from a high tumor-burden plasma sample (MEL-01, estimated tumor fraction of 22%) with aligned reads from a cancer-free control (CTRL-05). Reads were mixed to create 80x bam files harboring 10^−7^, 10^−6^, 10^−5^, 10^−4^, 10^−3^ or 10^−2^ tumor fraction. Tumor fractions were estimated either without any denoising, or with Ultima-specific denoising and MRD-EDGE integration. Z-scores were calculated as described above.

### UMI WGS data processing

FastQ reads were adapter and UMI trimmed using cutadapt (version 2.10). Trimmed reads were then aligned to the human genome (version hg38) using bwa mem (with parameters -K 100000000 -p -v3 -t 16 -Y). Trimmed UMI’s were added to the alignment files as an additional RX tag. Single-strand and duplex correction was carried out using the fgbio suite of tools (version 2.0). Because duplex correction via fgbio requires paired-end reads, we created a synthetic R2 read directly from single-end bam alignment files. R2 reads were built using the same mapping information (such as CIGAR string and mapping quality) and read information (such as sequence and qualities) as the R1 read. The subsequent paired-end alignment file was grouped by UMI (fgbio GroupReadsByUMI with parameters -m0 -s paired -e 1). Single-strand consensus sequences were created with CallMolecularConsensusReads and duplex consensus sequences were created using CallDuplexConsensusReads. Consensus reads were filtered with FilterConsensusReads and remapped to the human genome. Single-strand and/or duplex metrics (such as consensus read depth, consensus error rate, number of Ns on the consensus molecule, number of reads with matching UMIs) and mapping information were integrated as additional read tags to the original single-end alignment file. Variant frequencies in the original alignment files (without denoising) were calculated using lofreq^67^ (version 2.1.3a, with all filtering modes disabled). The original single-end bam, with the additional single-strand or duplex tags, was processed through the FlowFeatureMapper tool, (described above) which allows for the processing of the additional UMI tags, to obtain putative variants. Variants in duplex-resolved data were filtered based on the following conditions: <u>i</u>) duplex read must contain less than 4 Ns; ii) variant must not be within 25bp of either the 5’ or 3’ end of the molecule; iii) variant must be present on top and bottom strands; iv) all reads with the same UMI must have the 5bp flanking the variant match the reference. For single-strand correction data, variants were only considered if: <u>i)</u> the consensus read contains less than 4 Ns; ii) the variant must not be within 25bp of either the 5’ or 3’ end of the molecule; iii) all reads with the same strand-specific UMI must have the 5bp flanking the variant match the reference. Finally, UMI-agnostic denoising was performed by filtering by: i) variant position in read (the variant cannot be within 25bp of an end of the read); ii) template length (must be lower than 200bp); iii) mapping quality (cannot be below 60); iv) edit distance (must be below 4) and v) total variants on the read (must be below 11).

### Duplex, single-strand, and UMI-agnostic error rates in mouse PDX plasma samples

Denoising was performed as described above. Variants at a given genomic position, for each correction method, were compared to the frequency of that variant in uncorrected datasets. If the variant occurred two or less times in an uncorrected dataset, the variant was considered an error. The error rate was defined as the sum of the errors divided by the total number of base pairs for that correction method. For example, the error rate for duplex datasets corresponded to the number of errors divided by the total number of mapped base pairs from consensus duplex reads.

### Duplex, single-strand, and UMI-agnostic error rates in human plasma samples

We used samples from patient MEL-12 (n = 5 plasma samples) to estimate error rates. Here, we benefited from n = 5 Ultima-sequenced datasets, n = 5 Illumina-sequenced datasets and n = 1 WGS of normal DNA (PBMC, Illumina sequencing). Variants were identified in all datasets using lofreq^67^ (without filtering) or HaplotypeCaller^60^ (using GATK best standard practices) for plasma and PBMC data, respectively. For each Ultima-sequenced plasma sample, a patient-specific SNV mask was created by tabulating all variants appearing in any of the other 10 datasets, thereby extensively removing somatic and germline mutations specific to MEL-12. Variants were intersected with the mutation profiles of the 107 melanoma tumor datasets obtained from PCAWG (variants from the SNV mask were ignored). The error rate per PCAWG tumor was defined as the number of variants in the sample matching tumor mutations, divided by the number of tumor mutations multiplied by the average depth of sequencing of the sample.

### Statistical analysis

Statistical analysis was performed in R (version 3.6). Boxplots were generated using the ggplot2 R package. The lower and upper ends of the boxes represent the 25^th^ and 75^th^ percentiles of the data, respectively, and the horizontal line represents the median. The whiskers represent at most 1.5 times the IQR.

---

**Supplementary Figure 1.**
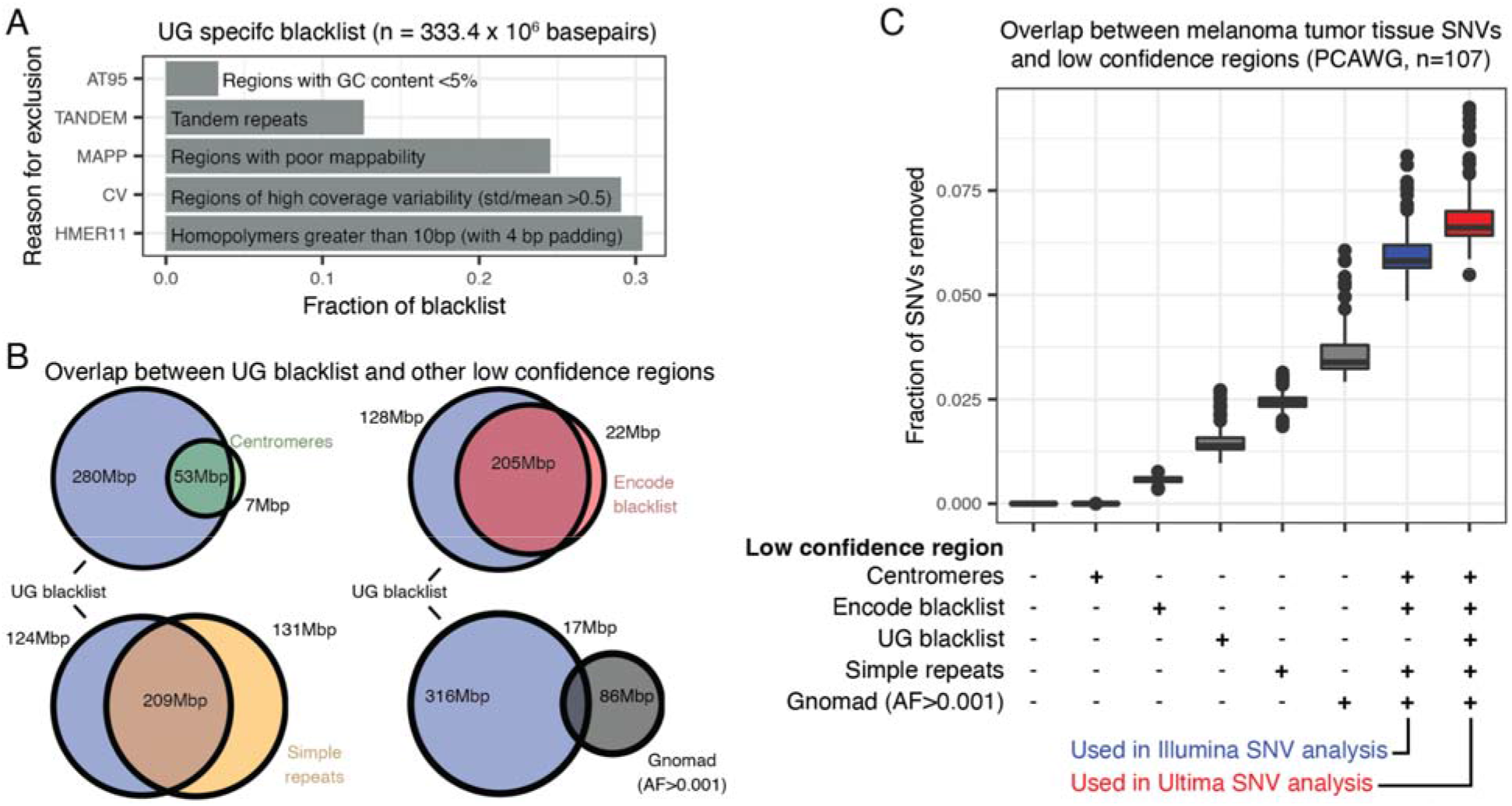
Effect of artifact blacklisting on single nucleotide variant detection. A. Ultima Genomics (UG) specific blacklist includes regions with low GC content, tandem repeats, regions with poor mappability, regions with high coverage variability and regions with homopolymers greater than 10 base pairs. B. Overlap between UG specific blacklist and other publicly available low confidence regions C. Effect of blacklists on the recovery of somatic single nucleotide variants (SNVs) in 107 melanoma tissue samples obtained from the Pan Cancer Analysis of Whole Genomes (PCAWG) consortium.

**Supplementary Figure 2.**
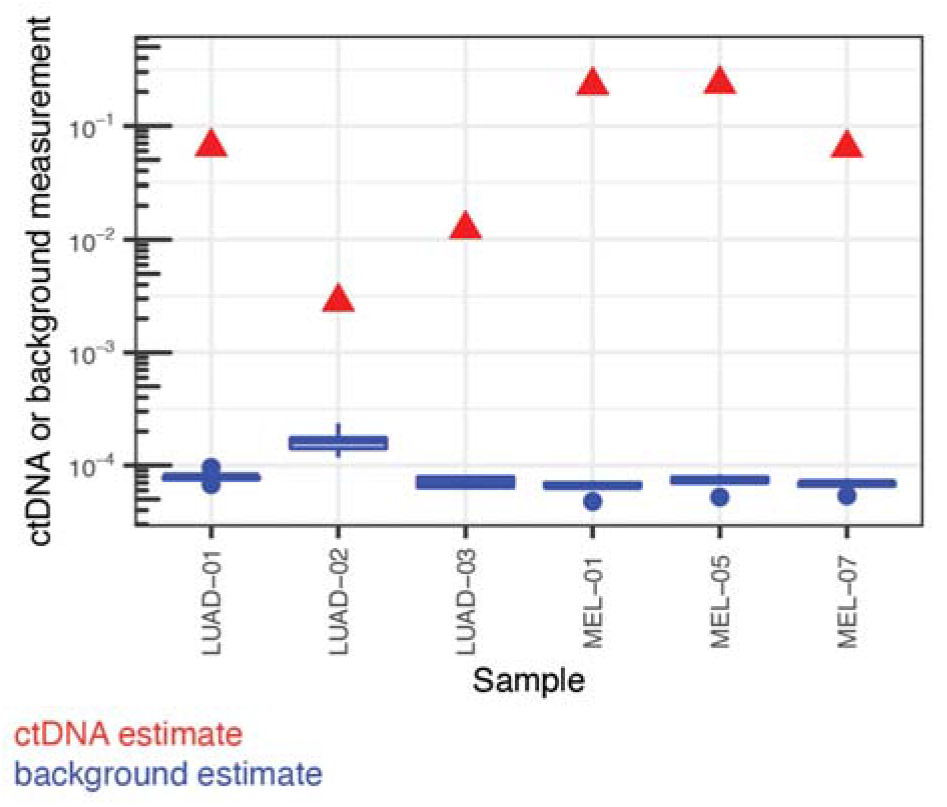
Estimated ctDNA (red) and error rates (blue) for Ultima whole genome sequencing samples. ctDNA estimates were obtained by intersecting putative plasma variants with the tumor-specific SNVs. Background estimates were obtained by intersecting cancer-free samples (CTRL-01 to CTRL-05) with the tumor-specific SNVs, and couting the rate of false positives.

**Supplementary Figure 3.**
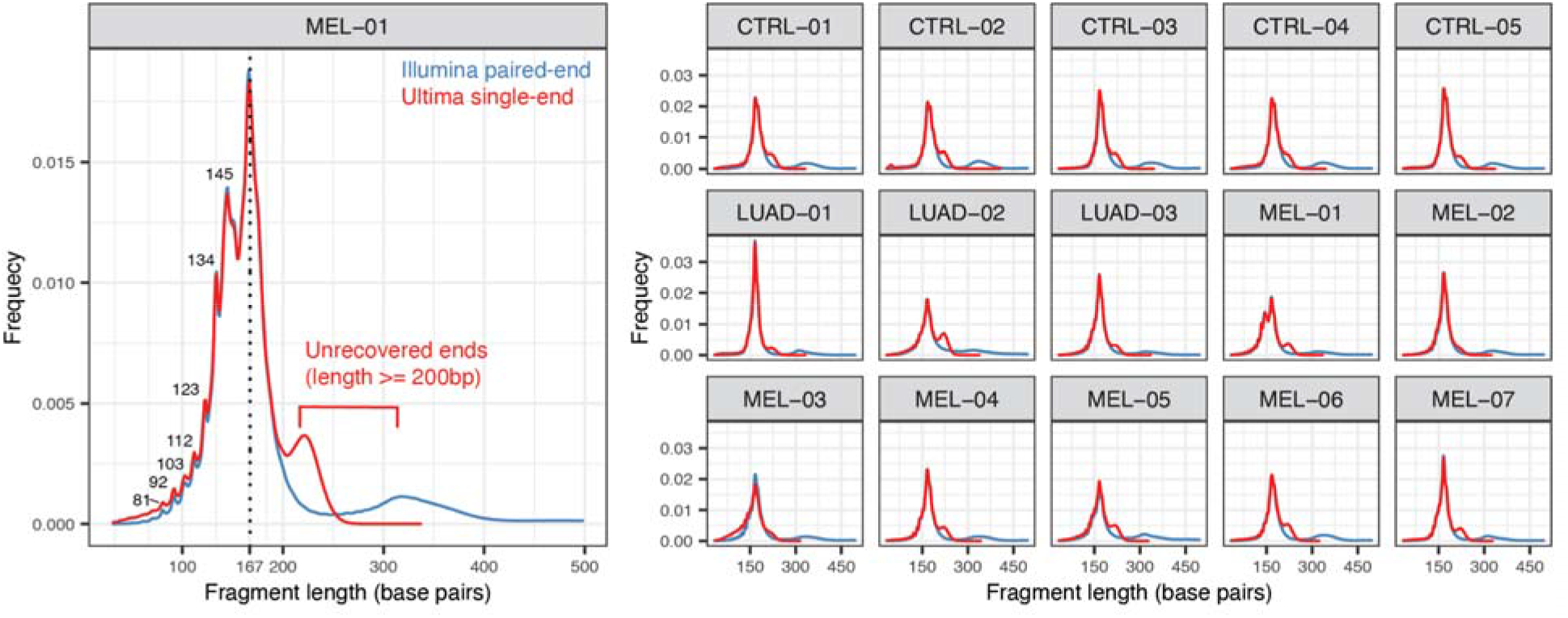
Cell-free DNA fragment lengths in single-end Ultima sequencing datasets matched with paired-end Illumina sequencing. Fragment lengths are accurately recovered between single-end Ultima reads when compared to paired-end Illumina sequencing for cell-free DNA molecules below 200 base pairs.

**Supplementary Figure 4.**
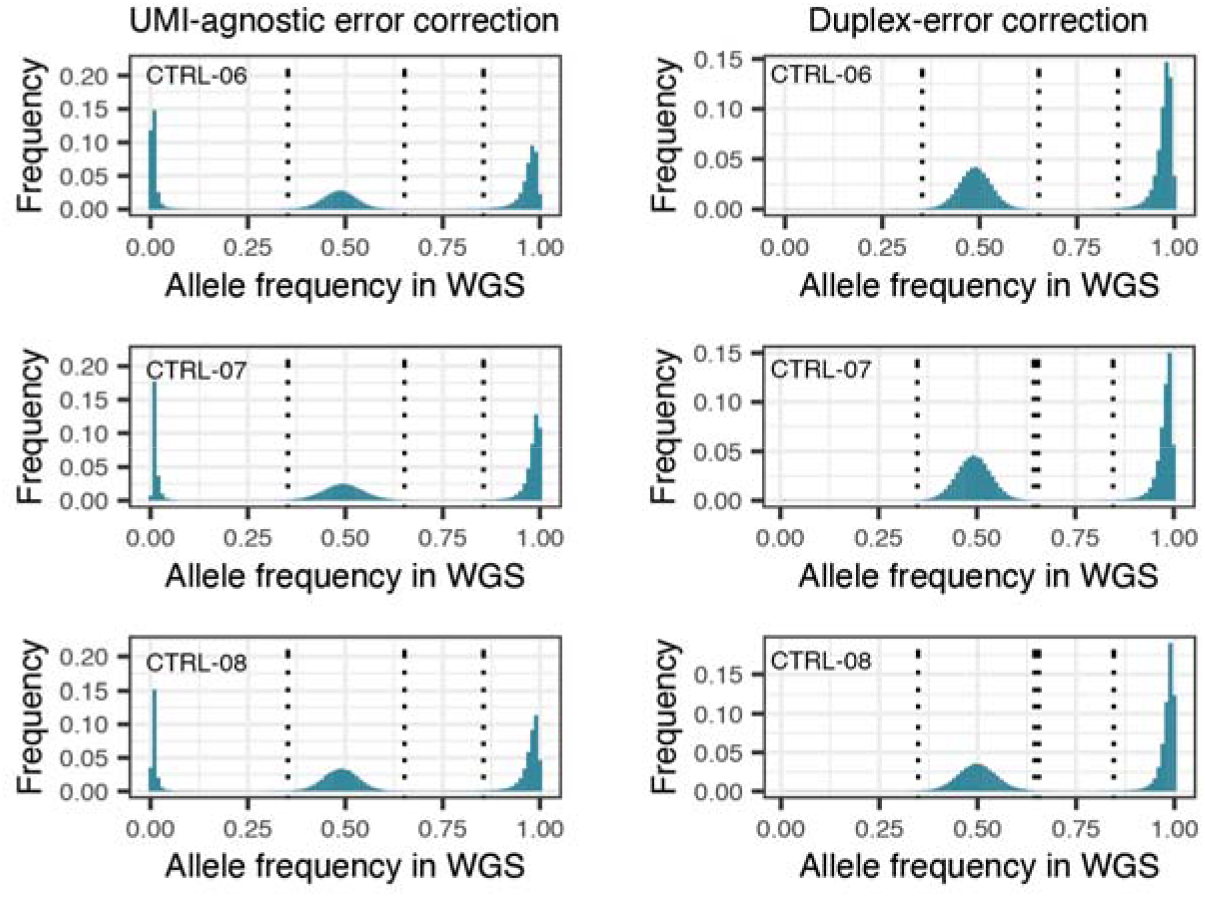
Variant allele frequencies (calculated using unfiltered sequencing reads) in positions where a variant was found using UMI-agnostic denoised reads (left column) and in duplex corrected reads (right column).

**Supplementary Figure 5.**
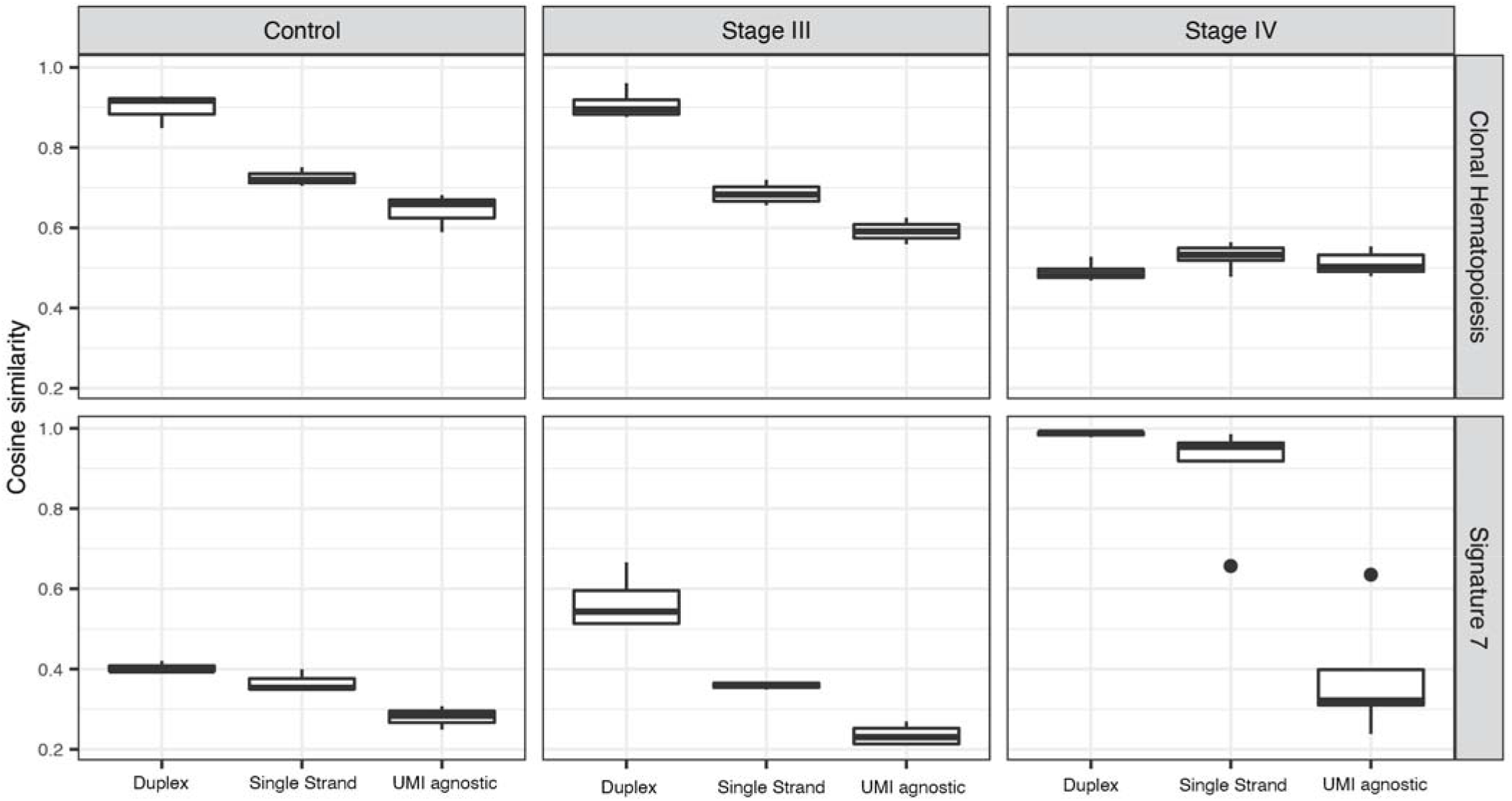
Cosine similarities in high burden and cancer-free samples for UV and CH-associated signatures and SBS7 and CH, respectively.

**Supplementary Figure 6.**
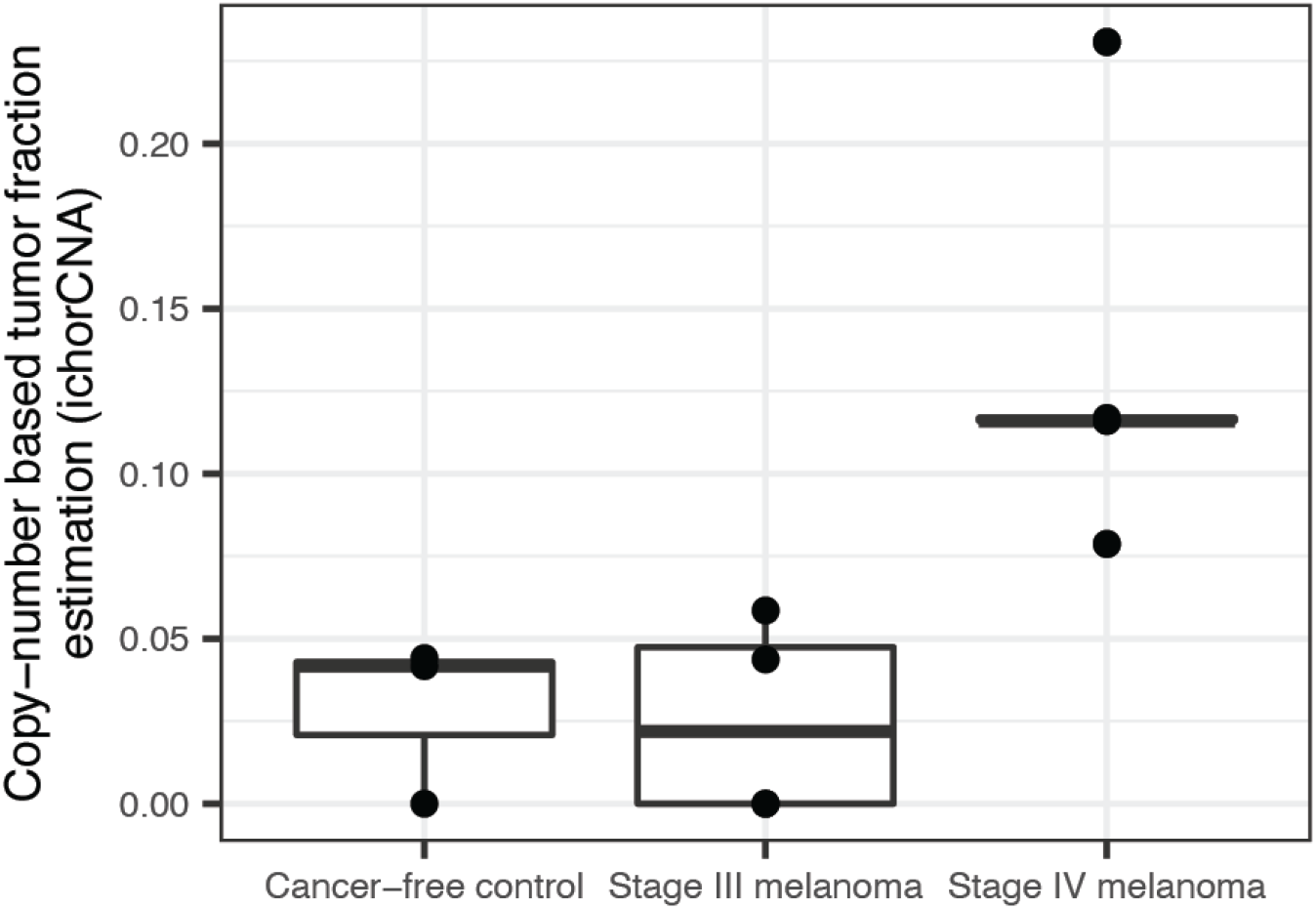
Tumor-agnostic copy-number based tumor fraction estimation in cancer-free control samples (n=3) and pre-surgery melanoma plasma (n=4).

